# Secretory vesicle trafficking in awake and anesthetized mice: differential speeds in axons versus synapses

**DOI:** 10.1101/268078

**Authors:** Johannes Knabbe, Joris Nassal, Matthijs Verhage, Thomas Kuner

**Author notes:** Equal contribution (Johannes Knabbe, Joris Nassal). Corresponding author: Thomas Kuner.

## Abstract

Neuronal dense core vesicles (DCVs) transport many cargo molecules like neuropeptides and neurotrophins to their release sites in dendrites or axons. The transport properties of DCVs in axons of the intact mammalian brain are unknown. We used viral expression of a DCV cargo reporter (NPY-Venus/Cherry) in the thalamus and two-photon *in vivo* imaging to visualize axonal DCV trafficking in thalamo-cortical projections of anesthetized and awake mice. We found an average speed of 1 μm/s, maximal speeds of up to 5 μm/s and a pausing fraction of ^~^11%. Directionality of transport differed between anesthetized and awake mice. In vivo microtubule +-end extension imaging using Macf18-GFP revealed microtubular growth at 0.12 μm/s and provided positive identification of antero- and retrograde axonal transport. Consistent with previous reports, anterograde transport was faster (^~^2.1 μm/s) than retrograde transport (^~^1.4 μm/s). In summary, DCVs are transported with faster maximal speeds and lower pausing fraction *in vivo* compared to previous results obtained *in vitro*. Finally, we found that DCVs slowed down upon presynaptic bouton approach. We propose that this mechanism promotes synaptic localization and cargo release.

**Key points:**

- Despite their immense physiological and pathophysiological importance, we know very little about the biology of dense core vesicle (DCV) trafficking in the intact mammalian brain.
- DCVs are transported at similar average speeds in the anesthetized and awake mouse brain compared to neurons in culture, yet maximal speed and pausing fraction of transport were higher.
- Microtubule +-end extension imaging visualized microtubular growth at 0.12 μm/s and revealed that DCVs were transported faster in the anterograde direction.
- DCV transport slowed down upon presynaptic bouton approach, possibly promoting synaptic localization and cargo release.
- Our work provides a basis to extrapolate DCV transport properties determined in cultured neurons to the intact mouse brain and reveal novel features such as slowing upon bouton approach and brain state-dependent trafficking directionality.

## Introduction

Neuronal dense core vesicles (DCVs) represent a diverse group of large secretory granules containing an electron-dense core as defined by electron microscopy. DCV cargo molecules differ between neuronal subtypes and include a variety of signaling molecules like neuropeptides, neurotrophic factors (such as BDNF), guidance cues, extracellular proteases and others. These signaling molecules are important for a variety of functions including synaptic plasticity, neurotransmission, neuronal development and survival^1–4^. Moreover, DCV cargos play important roles in several diseases such as social anxiety disorders (oxytocin and vasopressin^5^), stress disorders (NPY)^6^ and neurodegenerative diseases such as Alzheimer’s disease^6^. Despite their immense physiological and pathophysiological importance, we know very little about the biology of DCV trafficking in the intact mammalian brain.

DCVs are produced in the Golgi network^7^ and transported via microtubule-dependent motor proteins^8–10^ towards their axonal or dendritic release sites. In contrast to synaptic vesicles, the content released by DCVs is not recycled locally, but degraded by membrane- or circulating metallo-endopeptidases^11^. The highly dynamic transport of DCVs^12–17^ ensures a reliable supply to often very distant locations in the axon and dendrites. Previous studies have shown an anterograde and retrograde cycling of DCVs through the axon with sporadic pauses in axonal *en-passant* boutons, also referred to as varicosities, which was concluded to represent an activity-dependent capture mechanism^13,18^. The function of this mechanism is probably to enhance availability of DCVs at synapses to facilitate regulated secretion. The fusion of DCVs with the plasma membrane mostly occurs at/near synapses and can be induced with electrical high-frequency stimulation *in vitro*^14,16,19^. To date, it is not known, if DCV transport *in vivo* is influenced by properties of the surrounding neuropil. Studies addressing DCV trafficking in the intact mammalian brain are lacking. The dense packaging of axonal and dendritic compartments within the neuropil could mechanically affect local trafficking conditions, in particular when considering that axons can have diameters of less than 100 nm^20^, about the size of one DCV. Furthermore, neuro-glial interaction could affect DCV trafficking. In addition to these factors, different functional states of the brain such as sleep or wakefulness may have an impact on the properties of DCV trafficking and release. On a cellular level, different patterns of spontaneous and evoked activity in neurons in the intact brain, known to be more extensive than typically present in cultured neurons or acute brain slices, could significantly affect DCV transport properties. Until now, DCV transport was only studied in neuronal cell culture and model organisms like *D. melanogaster*^13^ and *C. elegans*^8^. Hence, to understand the contribution of aforementioned factors to DCV trafficking and to extrapolate published data obtained in reduced cell culture or brain slice preparations to living and behaving mammals, DCVs need to be imaged within the intact brain.

We established an *in vivo* DCV imaging approach using chronically implanted cranial windows and two-photon microscopy in anesthetized and awake mice to describe the basic DCV transport characteristics in thalamo-cortical axons ramifying in the molecular layer of the primary motor cortex. We show that DCVs travel through thalamo-cortical axons *in vivo* with an average speed of 1 μm/s and maximal speeds of up to 5 μm/s with infrequent pauses. Furthermore, we show that DCVs preferentially slow down near presynaptic *en-passant* boutons. We propose this mechanism helps to promote synaptic localization and release of neuropeptides and neurotrophins.

## Methods

### Ethical approval

All experiments were conducted in accordance with the German animal welfare guidelines and approved by the Regierungspräsidium Karlsruhe. The authors declare that the reported experiments comply with the journals ethical principles and their checklist on animal experimentation. Male C57Bl/6N mice were acquired from Charles River Laboratories (Sulzfeld, Germany) at an age of 8 weeks. The mice were kept at a 12/12-hour dark/light cycle synchronized with the local day-night cycle. Water and food were available ad libitum, except for the imaging sessions. Mice were generally housed in ventilated racks with up to 3 animals per cage, but were separated after surgery to minimize the risk of injury.

### Immunohistochemistry

For immunohistochemistry, mice were transcardially perfused with 4% paraformaldehyde (PFA) in phosphate-buffered saline (PBS). The brains were post fixed overnight in 4 % PFA and then sliced into 70 μm thick sections and stored in PBS. For increased permeabilization, the slices on which the Chromogranin stains were performed on, were treated with 1% sodium dodecyl sulfate (SDS) in PBS for 5 minutes before the antibody stainings. For antibody staining the slices were incubated for 1 hour in PBS containing 5% normal goat Serum (NGS), 1% bovine serum albumin (BSA), 1% cold fish gelatin and 0.5% Triton X-100. Incubations with primary antibodies were done overnight at 4°C. The secondary antibodies were incubated for 1-2 hours at room temperature. Primary antibodies were: Bassoon (Enzo Life Sciences, SAP7F407; 1:1000), Chromogranin A (SySy, 259003; 1:500), Chromogranin B (SySy, 259103, 1:500), Early endosomal antigene 1 (Eea1) (Abcam, ab2900; 1:200), Homer1 (SySy, 160003; 1:200), Lamp-2 (Santa Cruz Biotechnology, sc-18822; 1:200, kind gift from Ralph Nawrotzki), LC-3 (Medical & Biological Laboratories, M-152-3; 1:100; kind gift from Ralph Nawrotzki), Piccolo (SySy, 142104; 1:1000), Rab5 (Abcam, ab18211; 1:200). Alexa Fluor antibodies from Invitrogen were used as secondary antibodies in a dilution of 1:500. The slices were mounted in SlowFade Gold (Life Technologies) and imaged on a Leica SP8 inverted confocal microscope with a 63x oil immersion Objective (NA = 1.4) and maximal resolution in xy and z (0.08 μm × 0.08 μm × 0.3 μm).

### Genetic labeling of DCVs and thalamaco-cortical axons using viral gene transfer

DCV were labeled in vivo in mice by viral expression of a fusion protein consisting of Neuropeptide Y (NPY) and the yellow fluorescent protein Venus (NPY-Venus, see SFig. 1) or the red fluorescent protein mCherry^14,15^, driven by the synapsin promoter. Axonal projections were labeled by expressing mCherry under the control of the CAG (chicken-beta-actin) promoter in thalamic projection neurons. Microtubule plus-ends were marked by expression of GFP fused to the first 18 N-terminal amino acid residues of the microtubule plus-end marking protein Macf43 fused to the two-stranded leucine zipper coiled-coil sequence corresponding to GCN4-p1 under control of the synapsin promoter as described elsewhere^24^ (GFP-GCN4-MACF18).

Recombinant adeno-associated virus (AAV) particles of the chimeric 1/2 serotype were produced with the plasmids described above^37^.

### Craniectomy, stereotaxic injection and chronic window implantation

For viral injection^38^ and craniectomy (protocol adapted from Holtmaat et al. 2009^21^), mice were anesthetized with an intraperitoneal (i.p.) injection of a mixture of 0.48 μl fentanyl (1 mg/ml; Janssen), 1.91 μl midazolam (5 mg/ml; Hameln) and 0.74 μl medetomidin (1 mg/ml; Pfizer) per gram body weight each and placed in a stereotaxic headholder (Kopf). Before cutting of the skin, xylocain solution (1%; Astra Zeneca) was injected subcutaneously. A small hole in the skull was made with a dental drill over the injection site. Then, 1 μl of a 1:1 mixture of the NPY-Venus and mCherry or a mixture of MacF-GFP and NPY-mCherry AAV particles at similar infectious titers was slowly injected into the right medio-dorsal thalamus (Coordinates from bregma: x = 1.13, y = −0.82, z = −3.28). Afterwards a circular craniectomy (approx. 6 mm, center positioned 1 mm right of bregma) was drawn with a dental drill. The dura was carefully removed. A sterile round #0 coverslip with a diameter of 6mm (cranial window) and a custom made round plastic holder surrounding it for head fixation were cemented on the skull with dental acrylic (Hager & Werken).

After surgery, mice received i.p. a mixture of 1.86 μl naloxon (0.4 mg/ml; Inresa), 0.31 μl flumazenil (0.1 mg/ml; Fresenius Kabi), 0.31 μl antipamezole (5 mg/ml; Pfizer) in 3.72 μl Saline (0.9%; Braun) each per gram body weight to antagonize the anesthesia. Mice were given carprofen (Rimadyl, 5 mg/kg; Pfizer) in saline before surgery and during the following days in intervals of 12 hours for 3 days. Mice were single housed after surgery and typically imaged at least 21 days after the surgery. This waiting period is essential for the glial reaction below the window to subside^21^.

### *In-vivo* two-photon imaging

Two-photon imaging^39^ was performed with a TriM Scope II microscope (LaVision BioTec GmbH) equipped with a pulsed Ti:Sapphire laser (Chameleon; Coherent). 960nm was used for simultaneous excitation of both Venus and mCherry. For the experiments with MacF-Gfp and NPY-mCherry, a wavelength of 995nm was used. Imaging was performed with a 25X water immersion objective (Nikon MRD77225, NA = 1.1) and appropriate filter sets (Venus: 535/70 nm and mCherry: 650/100; Chroma). Fluorescence emission was detected with low-noise high-sensitivity photomultiplier tubes (PMTs, H7422-40-LV 5M; Hamamatsu). For the imaging sessions, anesthesia was induced with 6% isoflurane (Baxter) in oxygen and maintained at 0.8-1%. The anesthetized mouse was head fixed by clamping the plastic holder cemented to the skull. For awake imaging, the mouse was fixed over a rotatable disc treadmill (Figure 1; B) under isoflurane anesthesia and the isoflurane was shut off after fixing. The awake imaging was started at least 5 minutes after the end of isoflurane anesthesia. Frames were typically taken from an area of 196.6 × 196.6 μm at 1024 × 1024 pixel resolution with a frame rate of 0.942 Hz for 10 minutes per dataset using the galvanometer scanner of the TriM Scope II. Anesthetized imaging sessions lasted no longer than 1 hour and awake imaging sessions no longer than 20 min.

**Figure 1:**
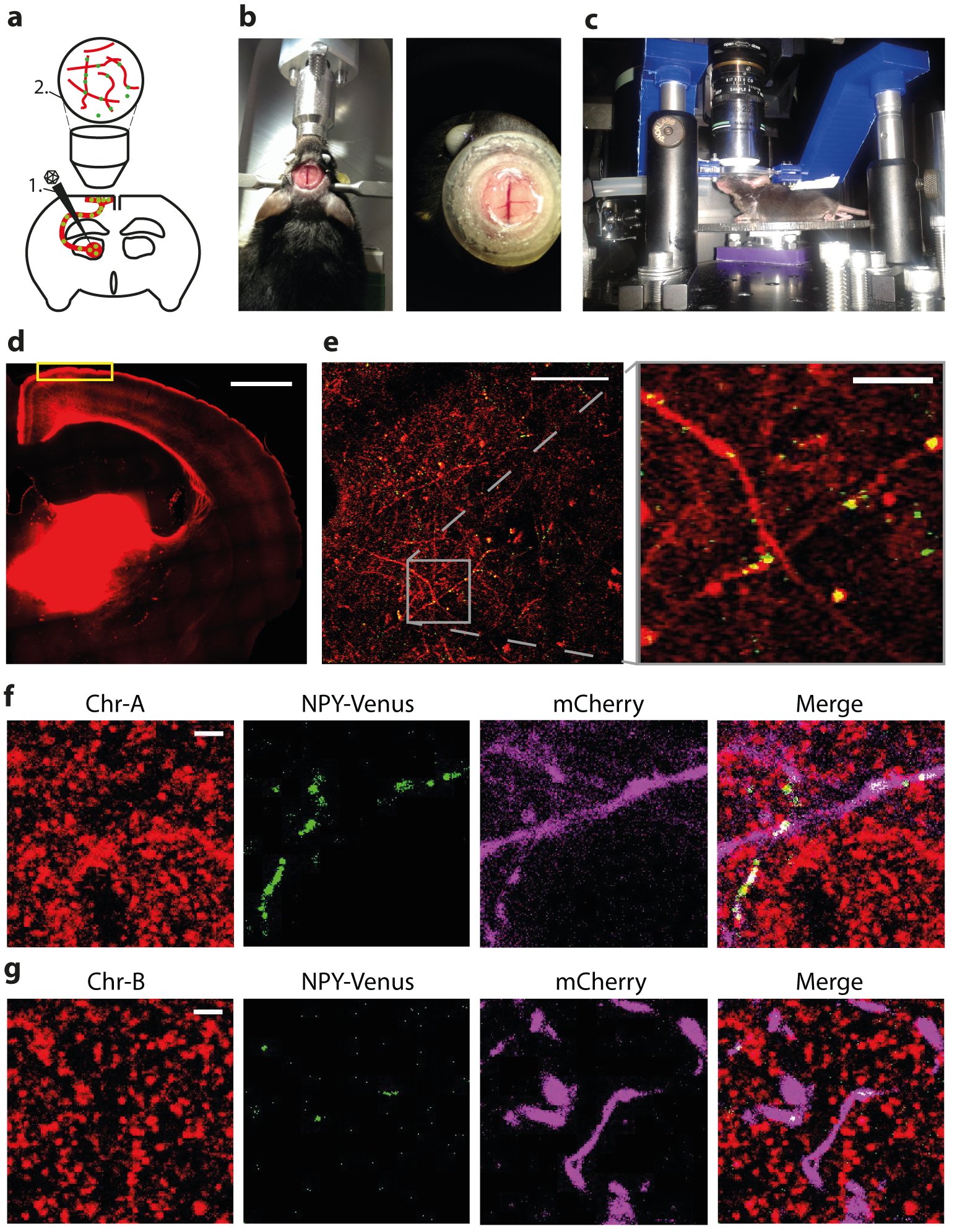
Experimental design and verification of DCV identity. (a) Injection of AAVs encoding NPY-Venus and mCherry into the area of the right medio-dorsal thalamus (1). Position of objective for imaging (2). Note the orientation of superficial axons in parallel to the brain surface. (b) Chronic window implantation after stereotaxic delivery. (c) Setup for awake imaging. The mouse is fixed via the crown, but able to move on a round freely rotatable disc. (d) Mosaic scan of thalamic injection site and axonal projection emanating from it (widefield epifluorescence imaging, scale bar = 1 mm). Yellow rectangle depicts region of interest imaged in primary motor cortex, 30-80 μm below the pial surface. (e) Example image of a single frame used for *in vivo* two-photon time-lapse imaging 40 μm below the pial surface. Scale bar 50 μm. Scale bar in the inset: 10 μm. (f) Average intensity projections of 5 consecutive confocal image planes showing an antibody staining against Chromogranin A (red), NPY-Venus (green) and mCherry-labeled axons (magenta). Scale bar 2 μm (g) as in (f) but antibody staining against Chromogranin B.

### Processing and analysis of imaging data

The acquired movies were registered for image movement in the xy-plane with the Fiji Plugin ‘Descriptor based series registration’^40^. Generally, during anesthetized imaging sessions, resulting displacement in xy was less than 10μm. The resulting stacks were deconvolved using the Richardson-Lucy algorithm implemented in Matlab (Mathworks) and a theoretical PSF, background subtracted and median-filtered. To only analyze moving particles, the average projection of the registered time lapse was subtracted from every image. For the semi-automated tracking of the DCV puncta the Fiji Plugin ‘Trackmate’^41^ was used. The automatically tracked parts were manually reviewed. The tracks were stopped when the puncta were not visible for more than two images. The resulting coordinates in x,y and t (time) were analyzed with custom written Matlab scripts. Only tracks exceeding 10 consecutive images were analyzed. The maximal speed per track was defined as highest mean of the speed out of 4 consecutive track points. The mean speed was averaged over all computed mean speeds of all tracks. The unidirectionality factor was defined as the fraction the DCV puncta moved in the direction of its general movement vector which was defined as the direction the vesicle moved from the first to the last track point.

The MacF18-GFP comets were analyzed using kymographs. Lines fitted over tracks in the kymographs were analyzed with an ImageJ macro written by Alessandro Moro which uses a kymograph plugin (Bioimaging and Optics Platform)^42^.

### Statistics

All statistics were calculated using Prism (GraphPad). Normality distribution of the data was tested using Kolmogorov-Smirnov, D’Agostino-Pearson and Shapiro-Wilk tests. Most analyzed data did not show a gaussian distribution. Hence, distributions of the datasets were compared with non-parametric two-sided Mann-Whitney or Kolmogorov-Smirnov tests. A p-value below 0.05 was considered statistically significant. Means reported in the text are given with the standard error of the mean (SEM) unless noted otherwise.

## Results

To image DCVs *in vivo*, vesicles and axons were fluorescently labeled with NPY-Venus, a previously optimized DCV reporter^19^ and mCherry, respectively, using viral gene transfer into the right medio-dorsal thalamus (Fig. 1a, SFig. 1). Thalamaocortical projection neurons were chosen as a model system, because their axons branch intracortically in the superficial layer 1 and run parallel to the surface of the brain, thereby providing ideal conditions for DCV imaging. Chronically implanted cranial windows (Fig. 1b) permitted *in vivo* two photon microscopy in anesthetized or awake mice (Fig. 1c)^21^. Three weeks after viral injection and chronic window insertion, imaging commenced using 960 nm excitation for simultaneous visualization of axons and labeled DCVs. The thalamic injection site and mCherry-labeled axonal projections reaching the surface of the brain are schematically indicated in Figure 1d, with the yellow rectangle indicating an area situated 30-80 μm below the pial surface, from which time-lapse data were acquired. An example of an *in vivo* two-photon imaging experiment is shown in Figure 1e, illustrating that labeling of thalamocortical axons with mCherry allowed detection of axons positioned within a single imaging frame, an important precondition to monitor moving DCVs (yellow dots in Fig. 1e, SMov. 1). The signal of analyzed vesicles shows a mean signal to noise ratio between 2:1 and 4:1 as measured by the mean fluorescence of the vesicle divided by the background, thus providing reliable imaging conditions.

The time-lapse movies regularly showed large static fluorescent clusters that did not overlap with the mCherry labeled axons. These might represent either cellular autofluorescent debris or released fluorescent cargo of the DCVs that has been taken up by cells, possibly astrocytes or microglia. Co-staining for an astrocytic marker and a marker of early endosomes demonstrated an overlap of a portion of this fluorescent cargo and astrocytic early endosomes (data not shown).

To control for the correct packaging of the DCV cargo reporter, NPY-Venus, into DCVs under *in vivo* conditions, we immunostained fixed brain slices with the endogenous DCV markers Chromogranin A and B (Chr-A, Chr-B). NPY-Venus-labeled DCVs were detected in mCherry-labeled axons and overlapped with Chr-A and Chr-B stainings (Fig. 1f,g). We found no overlapping fluorescent signals of NPY-Venus puncta with other cellular structures such as endosomes, lysosomes, autophagosomes or presynaptic proteins (SFig. 2). To exclude that expression of the DCV cargo reporter induced the formation of DCVs in cells that may not *a priori* produce DCVs, we labeled thalamic axons with mCherry and immunostained for Chr-A (SFig. 3). This experiment demonstrates the presence of DCVs in thalamocortical axons independent of the NPY-Venus expression. In summary, these results suggest that the DCV cargo reporter, NPY-Venus, correctly labeled DCVs in neurons that endogenously produce DCVs, consistent with previous observations made in cultured neurons^15,22^.

### Visualizing DCV movement in axonal projections *in vivo*

DCV trafficking in thalamocortical axons was monitored by time lapse imaging of DCV reporter fluorescence and axonal fluorescence in single focal planes at 1 Hz. Individual DCV puncta can be assigned to the axons they are traveling in (Fig. 2a) and subsequently tracked (Fig. 2b). Small fluctuations in the apparent size of the puncta along the trajectory may arise from axons not running perfectly in parallel to the imaging plane, so that different amounts of fluorescent proteins in the vesicles are excited (see SFig. 1). Alternatively, changes in the z focus position in response to slight movements of the tissue may account for fluctuations in apparent size of the puncta. The average length of axonal segments analyzed was 30 ± 10 μm and time-lapse movies lasted for typically 10 min. At t = 2 s, the DCVs marked yellow and blue in Fig 2b appear to travel within the same diffraction-limited focal volume. During the time period from 2 s to 14 s, the two moving puncta travel with different speeds (1.6 μm/s; 0.6 μm/s) in the same direction. This is further illustrated in the kymograph (Fig. 2c,d). Hence, tracing of labeled DCVs can be achieved with the approach introduced here, including quantitative analyses.

**Figure 2:**
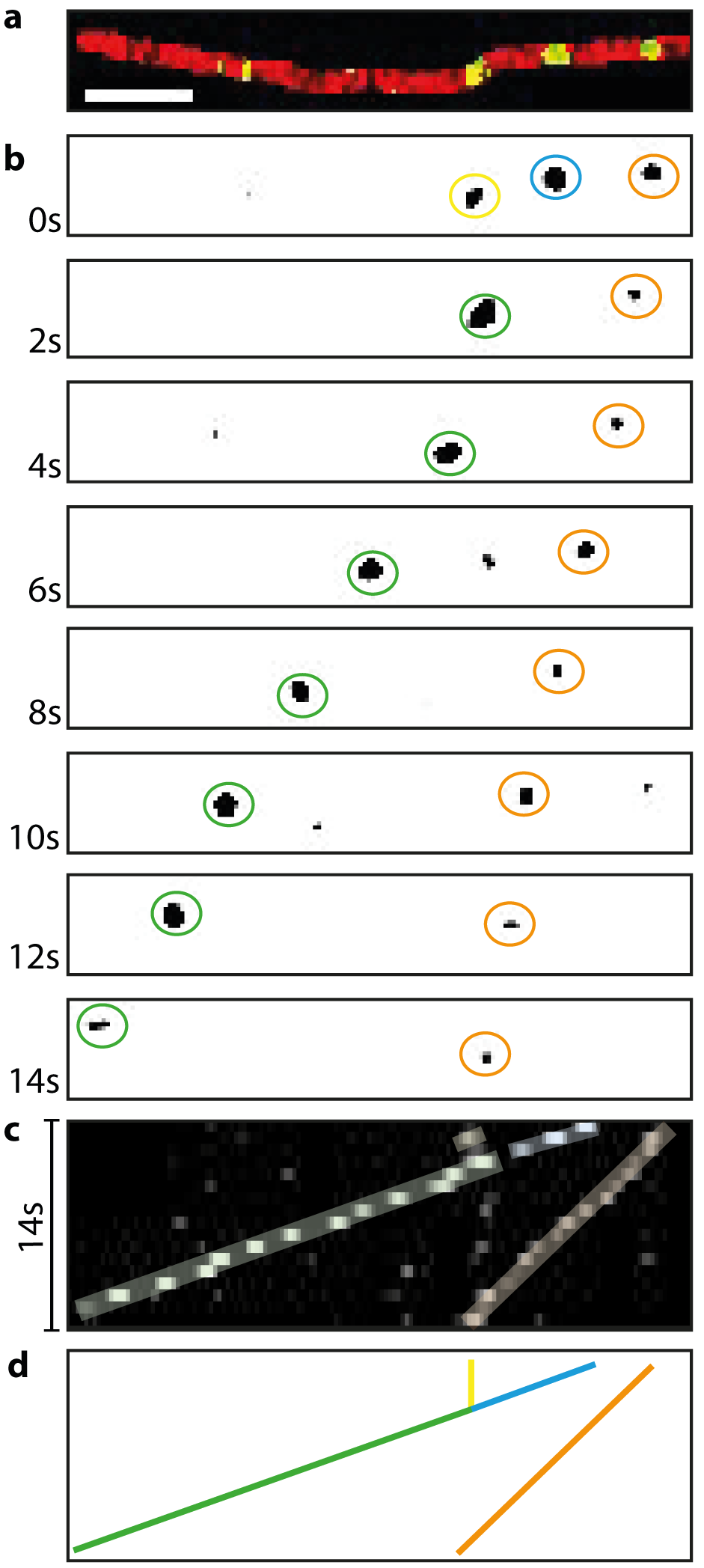
Time-lapse images and kymograph of moving DCVs in a single axon. (a) Single two-photon image frame showing an axon labeled with mCherry (red) and The DCV cargo reporter NPY-Venus (green). Scale bar 5 μm. (b) Time-lapse images of the green channel in panel a. DCVs labeled blue and yellow at t=0s travel together (green) in subsequent frames. (c) Kymograph of time- lapse in b. (d) Illustration of Kymograph for better visualization.

### Quantification of DCV trafficking characteristics *in vivo*

Following the experimental scheme shown in Figure 2, a total of 279 vesicles in 8 mice were tracked semi-automatically. The average speed of moving DCV puncta *in vivo* in anesthetized mice was 1.03 ± 0.03 μm/s (Fig. 3a, SFig. 4b). This is in agreement with previous observations in cultured mammalian neurons and with *in vivo* imaging performed in *D. melanogaster*, ranging from speeds of 0.75 μm/s to 1.25 μm/s for axonal DCV transport (see colored points in Fig. 3a)^9,12,23^.

**Figure 3:**
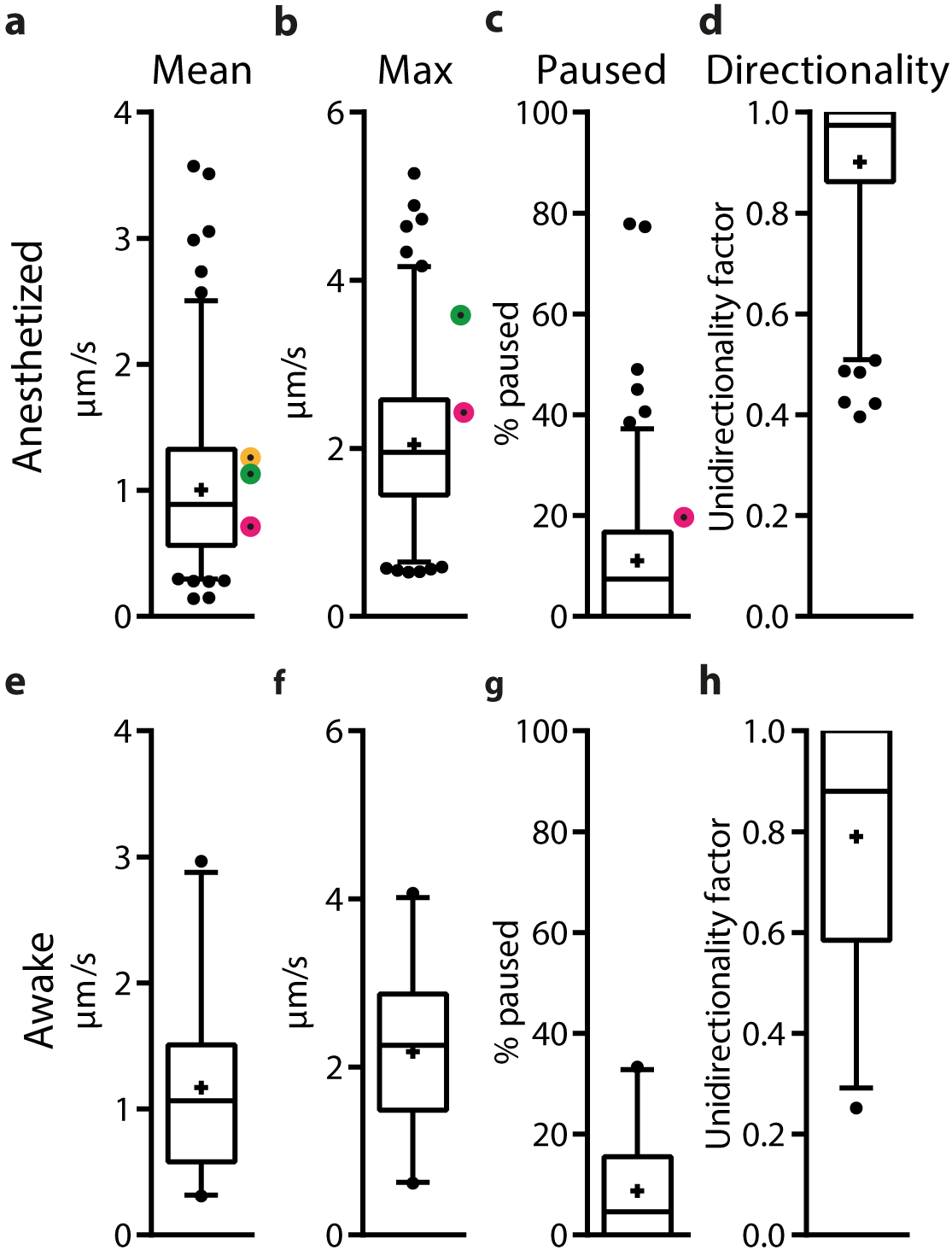
DCV movement dynamics resemble previous *in vitro* studies. (a-d) Boxplots of 279 tracked puncta taken from 8 anesthetized mice. Boxplots show the median and the 25 to 75 percentile, the whiskers show the 2.5 to 97.5 percentile. Individual points above or below the whiskers are outliers. Colored points represent previous reported data from in vitro studies. Magenta: de Wit et al. 2006, turquoise: Bittins et al. 2010, yellow: Kwinter et al. 2009. (b) Maximal speed averaged over 4 consecutive track points. (c) Pausing time as the percentage of images in which puncta moved with a speed below 0.2 μm/s. (d) Directionality of all vesicles. (e-h) Puncta tracked awake mice (n=46 puncta, 3 mice). No significant differences (Mann-Whitney test) were found for mean speed (p=0.12), max speed (p=0.28), and fraction paused (p=0.22), while the directionality differed highly significantly (p = 0.0001).

The maximum speed per track was defined as the highest average speed within 4 consecutive track points of each track. The average maximum speed of the tracked vesicles was 2.06 ± 0.05 μm/s (Fig. 3b), but we regularly observed speeds of up to 5 μm/s (*in vitro* data: 2.6 μm/s to 3.75 μm/s^12,23^).

DCV puncta often slowed down and stopped, but started moving again at a later time point (pausing). The pausing time was defined as the percentage of time that a punctum moved slower than 0.2 μm/s. This corresponds roughly to a displacement of one pixel per image. The overall pausing time of the tracked vesicles was 10.7 ± 0.7 % (Fig. 3c). This is lower than the 22% pausing fraction reported in cell culture^12^.

It was not possible to determine whether DCV puncta moved in the anterograde or retrograde direction within the axon, because the position of the soma could not be determined *in situ*. To analyze if vesicles moved back and forth, or if they followed a preferred movement vector, the directionality factor for the overall movement of a vesicle was calculated. This ‘unidirectionality factor’ was defined as the fraction of time the vesicle moved along its overall directional vector. The mean unidirectionality factor of all 279 analyzed tracks is 0.86 ± 0.01 (Figure 3d). This corresponds to an overall linear unidirectional movement of the puncta. The remaining 14 % of movements were in the opposite direction to the main movement vector.

To compare the amount of bidirectional and unidirectional movement in individual axons, only axons with more than 5 tracked puncta were investigated to avoid a possible bias towards unidirectional movement. In total, 25 axons were analyzed. The frequency distribution of the directionality in these axons shows that in approximately 40% of the axons DCVs move unidirectionally (SFig. 4a). Bidirectional movement was observed in approximately 60% of the axons.

To test possible confounding effects on the trafficking characteristics due to different expression time and thus different amounts of DCV cargo reporter expressed, two mice were imaged twice, at 2-3 weeks and 8-9 weeks after viral injection (SFig. 5). No significant differences in the transport speeds and other trafficking characteristics of the NPY-Venus puncta were found between these time points, suggesting that the amount of reporter expressed did not affect the trafficking properties of DCVs.

In summary, the basic DCV trafficking properties were similar in the anesthetized intact brain in comparison to a large set of trafficking data previously described in neuronal cell cultures. However, *in vivo*, the peak speed was higher and the pausing time lower.

### Comparison of DCV trafficking in anesthetized and awake mice

To control for potential influence of anesthesia, and thus brain activity levels in general, on DCV transport, three mice were imaged in the awake state. Minor axial movements exceeding 1-2 μm remove the axon stretch of interest from the imaging plane (see SFig. 1). Therefore, the base plate of the window was firmly fixed, both on the skull and in the holder, while placing the mouse on a rotatable disk (Fig. 1c). The axial focal plane remained stable, except when the mice started to run. These episodes were excluded from further analysis. 46 tracks in awake and resting mice were analyzed and trafficking characteristics from these tracks were compared to all tracks recorded from anesthetized mice. The mean speed of DCV puncta was 1.17 ± 0.1 μm/s in the awake state and did not differ from the mean speed of 1.03 ± 0.03 μm/s in anesthetized mice (Fig. 3e, SMov. 2). This is further illustrated by the large overlap of the frequency histograms of both distributions (SFig. 4b). The maximal speed and pausing time did not differ significantly between anesthetized and awake mice (Fig. 3f, g).

The only feature in DCV trafficking that was significantly different in the awake compared to anesthetized mice was the lower directionality factor of 0.79 ± 0.03 in awake animals compared to 0.9 ± 0.01 (MW, P= 0.0002) in anesthetized animals (Fig. 3h). This could potentially be explained by interfering movement of the animal that could not be corrected for during recordings. However, also intrinsic differences caused by the activity state of the brain may affect DCV movement directionality.

### Axonal microtubule dynamics *in-vivo*

To discriminate between anterograde and retrograde DCV trafficking *in vivo*, we co-expressed NPY-mCherry and MacF18-GFP, a minimal fragment of the microtubule-actin cross-linking factor 2 that gets selectively incorporated at the microtubule plus-end^24^. This allowed us not only to determine axonal orientation for trafficking analysis but also to characterize axonal microtubule growth dynamics. MacF18-GFP fluorescence accumulated at the growing microtubule plus ends referred to as comets (Fig. 4, SMov. 3). As overexpression of MT-plus end binding proteins can lead to the binding to all microtubules and not only the plus ends^29^, we analyzed these mice exclusively after 3-4 weeks of expression time, where punctate fluorescent and only faint cytoplasmic signals could be observed in axons. Analysis of the kymographs of Macf18-GFP comets allowed us to quantify movement speeds of axonal microtubules *in vivo*. 155 comets in 17 axons of 4 mice were analyzed. We found a mean speed of axonal microtubule polymerization *in vivo* of 0.12 ± 0.01 μm/s (Fig. 4d). This is slightly higher than the previously reported speed *in vivo* in dendrites^24^.

**Figure 4:**
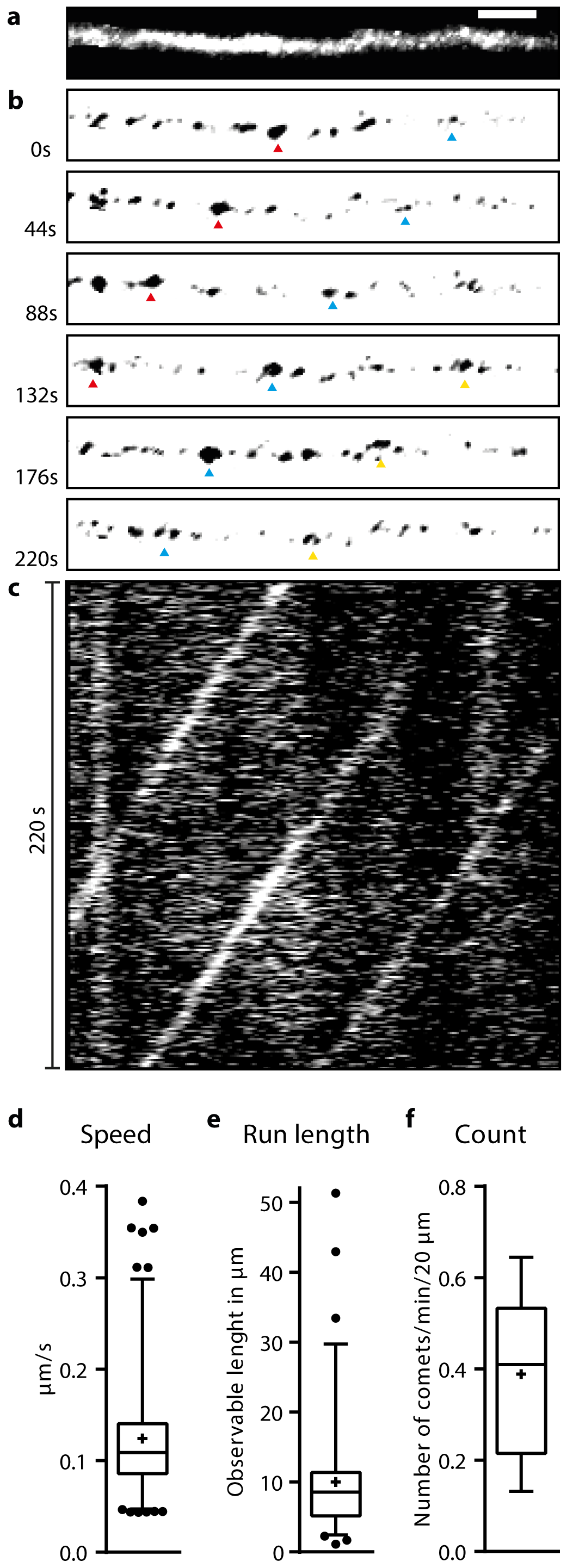
Axonal microtubule dynamics *in vivo*. (a) Maximum intensity projection of time-lapse of MacF18-GFP positive, manually segmented axonal stretch. Scalebar 5μm (b) Time-lapse of microtubule +end growth. Three comets are marked with colored triangles. (c) Kymograph of time-lapse shown in b. (d,e,f) Boxplots of mean speed, run length and number of comets per min per 20 μm.

The average run length of single comets was 10.02 ± 0.59 μm (Fig. 4e). To compare the number of comets per axon stretch length and time to previously reported numbers, we calculated the number of comets per 20 μm per min (Fig. 4f). With ^~^0.4 comets/min/20μm we observed fewer comets than the previously reported number of ^~^ 1.8 comets/min/20 μm^24^ in dendrites.

### Anterograde versus retrograde DCV trafficking

Assessing the run direction of MacF18-GFP puncta allowed us to define the directionality of DCV movement within the same axon as anterograde or retrograde with respect to the soma. The kymograph of an axon stretch in Figure 5a shows the unidirectional movement of multiple microtubule comets (green), thereby defining their movement direction as anterograde. The movements of multiple vesicles in the same axonal stretch (red) reveal the dynamic bidirectional transport of DCV.

**Figure 5:**
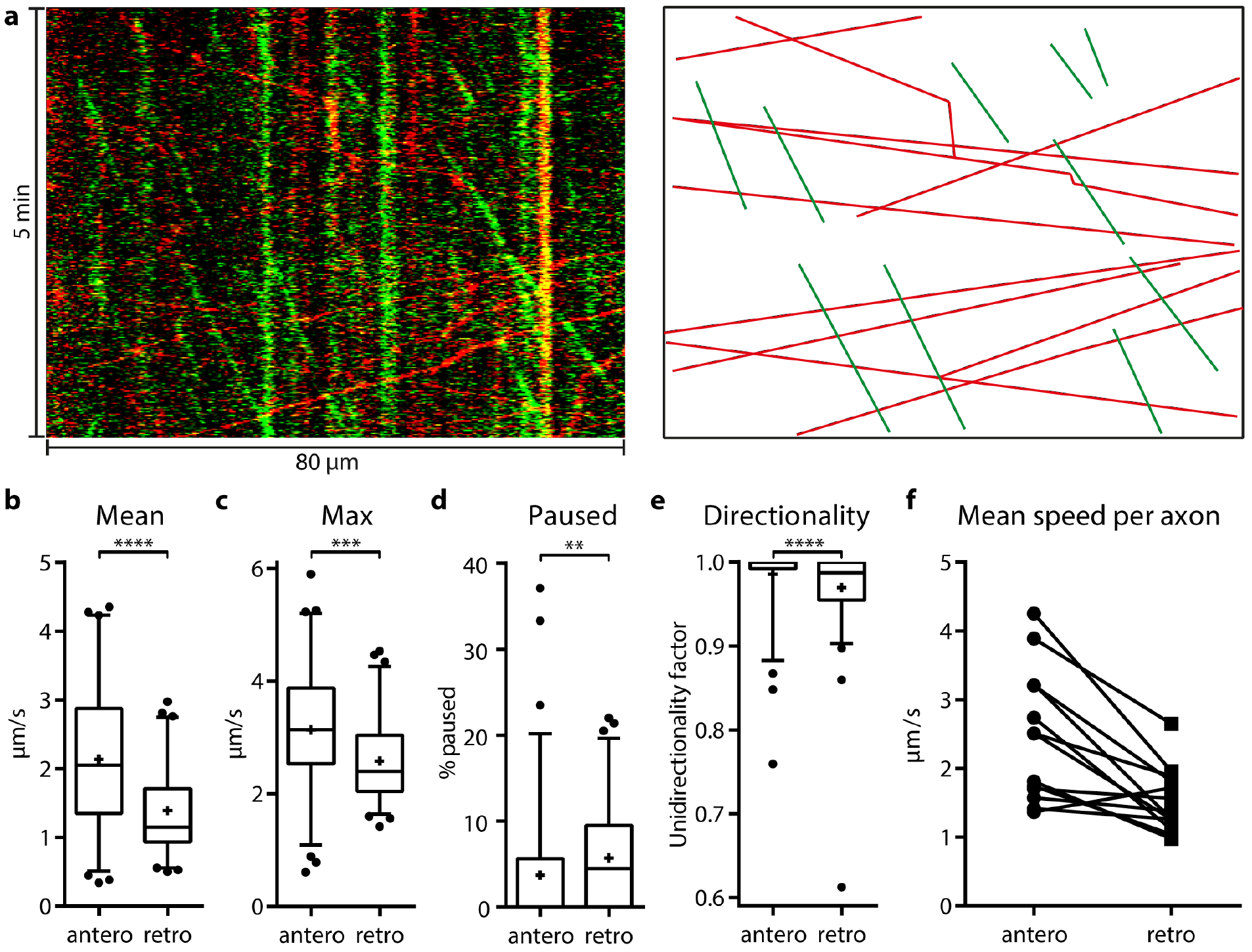
Anterograde versus retrograde trafficking of DCVs *in vivo*. (a) Left: Kymograph covering 5 min of an axon stretch with green puncta depicting MacF18- GFP comets and red puncta NPY-mCherry in DCVs. Right: Linear illustration of left kymograph. (b-e) Comparison of 78 tracked vesicles in anterograde and 67 tracked vesicles in retrograde direction in 3 different mice. MW test showed significant difference in mean and max speed as well as in pausing time and unidirectionality factor (p-values: <0.0001, 0.0002, 0.0021, <0.0001) (f) Comparison of mean speeds within axons showing more than 2 vesicles in each direction.

In the axons the mean speed of DCV transport in the anterograde direction is 2.14 ± 0.12 μm/s (n = 78) and in retrograde direction 1.4 ± 0.08 μm/s (n = 67; Fig. 5b). This analysis shows a significant difference in transport speed (Mann-Whitney-Test, p<0.0001). Additionally, we compared mean speeds of anterograde and retrograde transport within axons (with n > 2 vesicles in each directions). In almost all axons, the anterograde vesicles were faster than the retrograde vesicles (Fig. 5 f). The maximum speed, pausing time and unidirectionality factor show significant differences in the antero- and retrograde trafficking direction (Fig. 5 c,d,e). Thus, *in vivo* imaging revealed a difference in anterograde and retrograde trafficking speed comparable to that previously published in cultured neurons^12^ and invertebrates^8,25.^

### DCV trafficking is different near axonal boutons in vivo

While analyzing the tracking data, we noticed that some of the axons showed different trafficking features at sites that resembled *en-passant* boutons, visible as ellipsoid structures with a diameter of 2 - 4 μm (Fig. 6a). To confirm that these structures correspond to axonal boutons, we performed immunostainings for pre- and postsynaptic markers. Immunostaining against synaptophysin, a synaptic vesicle protein, and Homer1, a protein of the postsynaptic density, showed overlap of synaptophysin-positive puncta in ellipsoid structures in the axon (SFig. 6a) and revealed an enrichment of Homer1-positive puncta near putative boutons (SFig. 6b). These observations confirm that the ellipsoid structures of typical size and shape represent presynaptic boutons.

**Figure 6:**
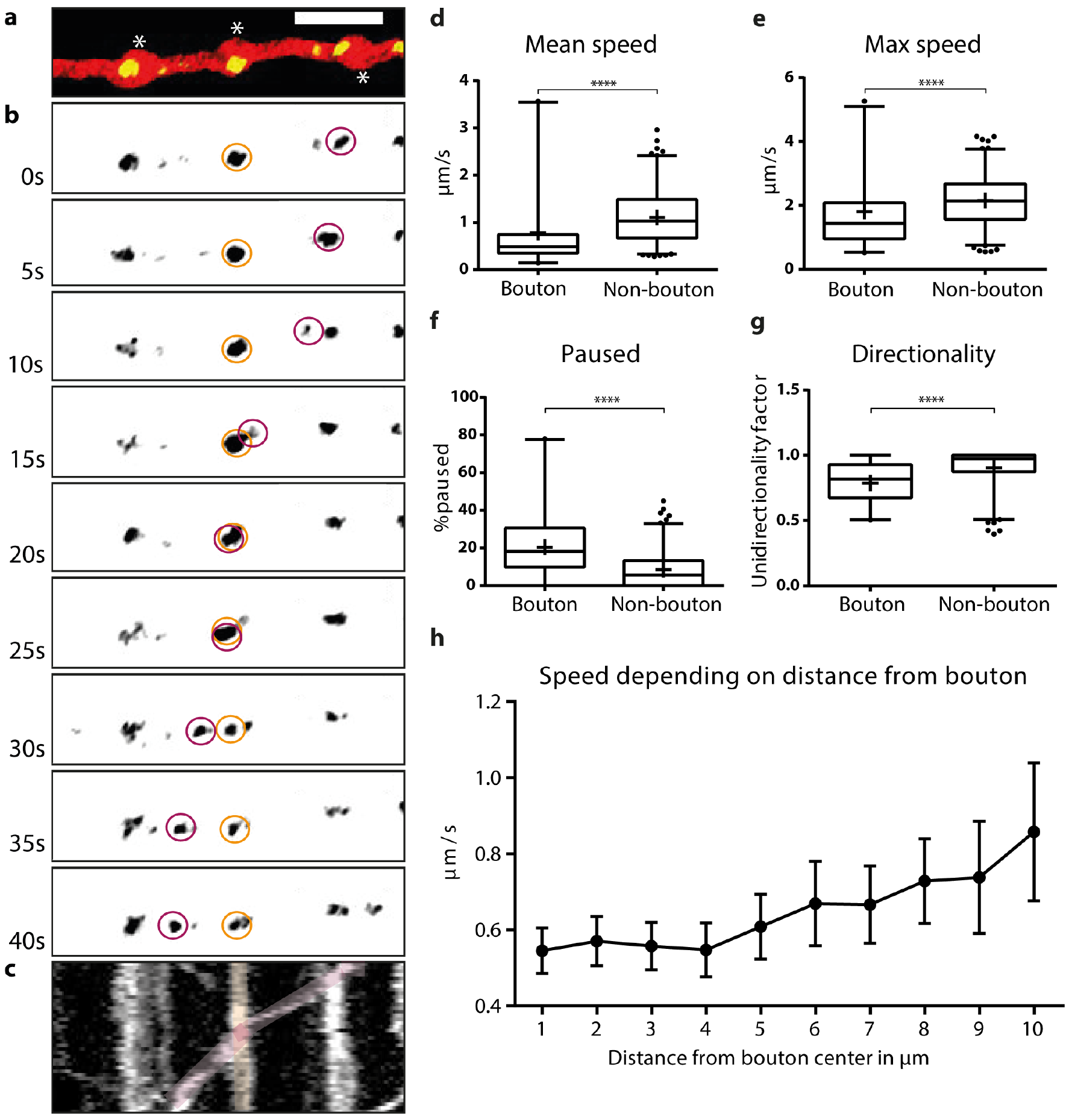
Reduced DCV trafficking speed at*en-passant* boutons. (a) Single two-photon frame showing a mCherry-labeled axon containing NPY-Venus puncta, white asterisks indicate *en-passant* boutons. (b) Time-lapse images of NPY-Venus puncta. Scale bar 5 μm. (c) Kymograph of time-lapse in b. (d-g) Comparison of 245 tracks from axons lacking boutons with 57 tracks from axons having visible boutons. The groups are significantly different in all analyzed trafficking characteristics (Mann-Whitney Test; p-values < 0.0001). (h) Mean speeds of tracks in relation to their distance to the bouton.

Figure 6a shows a representative stretch of axon harboring three boutons. Each of the boutons contains immobile (e.g. orange) and mobile puncta (Fig. 6b). Upon passing the bouton, mobile puncta slow down near the boutons (red, see also trajectory in kymograph, Fig. 6c). For the subsequent analysis, 57 tracks of axons with visible boutons from 3 mice were compared to tracks in axons lacking boutons. The average speed of DCV puncta was significantly lower in the axons containing boutons with a difference of 0.3 μm/s (Fig. 6d; bouton: 0.78 ± 0.11 μm/s; non-bouton: 1.10 ± 0.03 μm/s; MW: p < 0.0001). The maximal speed was significantly lower too, with a difference of 0.3 μm/s (Fig. 6e; bouton: 1.81 ± 0.16 μm/s; non-bouton: 2.16 ± 0.05 μm/s; MW: p < 0.0001). The pausing time of the vesicles was different with a higher fraction of pausing vesicles in/around boutons (Fig. 6f; bouton: 20.43 ± 2.14 %; non-bouton: 8.50 ± 0.61 %; MW: p < 0.0001). Finally, changes in movement directionality were increased in axons containing boutons (0.79 ± 0.02; non-bouton: 0.90 ± 0.01; MW: p < 0.0001).

To investigate further at which distance from the bouton center the reduction in speed occurred, all bouton center points were marked manually. Mean speeds of tracks outside a certain distance from the center were analyzed covering a range of 1 to 10 μm (Fig. 6h). A steady decrease in mean speed from ^~^ 0.9 μm/s to ^~^ 0.5 μm/s occurred within a distance of 10 μm towards the bouton center. We conclude that a lower mean speed of vesicles within and near boutons leads to longer residence of trafficking DCVs in presynaptic boutons, facilitating the recruitment of DCVs for secretion in synaptic boutons.

## Discussion

We characterized DCV trafficking dynamics in anesthetized and awake mice in axons of thalamaocortical projection neurons. Our experimental paradigm allows imaging of fluorescently labeled neuronal DCVs through chronic cranial windows in mice, transferring basic cell biology from the culture dish to the more complex and realistic environment of higher order systems. While many axonal DCV trafficking properties previously described *in vitro* are similar to those revealed here *in vivo*, we found a higher peak transport speed, a lower fraction of pausing DCVs and a slow down of DCV trafficking upon approach to axonal boutons. The latter is consistent with a preferential capture and release of DCVs at presynaptic sites in the anesthetized and awake mouse brain.

NPY-Venus-labeled punctate structures showed the trafficking features of DCVs and were confirmed as such using staining for Chromogranin A/B (Fig. 1f,g). Furthermore, NPY-Venus signals did not overlap with other (mobile) organelles such as endosomes, lysosomes or autophagosomes, or presynaptic proteins (SFig. 2). Together, these observations strongly indicate that NPY-Venus-labeled structures represent DCVs. The NPY-Venus puncta traveling through axons had different sizes and fluorescence intensity, suggesting that puncta may consist of either single or multiple DCVs (see SFig. 1), consistent with published *in vitro* data^14^. It has been argued that thalamo-cortical and pyramidal cortical neurons lack DCVs altogether^26^, although thalamic projection neurons have not been systematically investigated for DCVs. By selectively labeling thalamic projection neurons with mCherry and performing stainings of the endogenous DCV marker Chromogranin A, we could demonstrate that DCVs naturally occur in these neurons (SFig. 3). Hence, NPY-Venus- expression did not artificially induce DCV biogenesis and our study reports on physiologically relevant processes. In addition to axonal fluorescent clusters we also regularly found larger fluorescent clusters outside the axons. These clusters may reflect autofluorescent cellular debris near the pial surface, stable deposits of secreted DCV cargo or endocytosed fluorescent proteins within scavenger cells such as astrocytes or microglia, or in neurons. Such accumulations of DCV-cargo after their release is a well known phenomenon in coelomocytes of the nematode *C. elegans*^8^.

The DCV velocities and directionality of their transport determined here generally correspond well with previously reported data from *in vitro* experiments and studies in other model systems^8,9,12,23,25^. The higher peak speed and shorter pausing time observed *in vivo* might be due to different temperatures in cell culture experiments or the fact that axonal microtubule orientation is different in mammals and invertebrates^27^. Furthermore, neuronal activity patterns are probably different *in vitro* and *in vivo* and lead to a different amount of arrest, as DCV arrest is induced by activity^12^. Finally, we observed a wide range of DCV trafficking patterns: Faster vesicles passing slower ones in the same direction, vesicles first traveling alone, then together to finally be separated again, vesicles passing each other in different directions, most vesicles slowing down mid travel to pause for a short time and others traveling with a constant speed over the whole tracked distance. The frequent changes in movement speed and also directionality may be due to different motor proteins of the kinesin family bound to the same vesicle, possibly in combination with a dynein motor, thereby defining the speed of the vesicles as a product of the individual speeds of the motors involved^17^. These diverse *in vivo* trafficking patterns resemble patterns previously observed *in vitro*^12^. In conclusion, the general properties of DCV trafficking in the intact mouse brain are in agreement with previous findings in neuronal primary cell culture and invertebrates. This may be attributed to the high conservation of motor proteins that are responsible for DCV transport in all investigated model systems^8,25^.

Coexpression of the microtubule marker Macf18-GFP with the DCV marker NPY-mCherry allowed us to distinguish between anterogradely and retrogradely moving DCVs. The mean speed of all anterogradely traveling DCVs was 2.14 μm/s, approximately 50% faster than vesicles moving retrogradely, at 1.4 μm/s. This is in accordance with findings from cultured hippocampal neurons, but in opposition to findings at *Drosophila* neuromuscular junction, where DCVs move antero- and retrogradely at roughly the same speed^18,25^. Our findings are consistent with speeds reported for motor proteins recruited for axonal anterograde and retrograde movement, with kinesins moving DCVs considerably faster than dynein^28^. The overall higher mean speed reported in Figure 5 compared with that reported in Figure 3 might be attributed to the fact that we only analyzed axonal stretches containing multiple moving DCVs, potentially interfering with trafficking speed. Furthermore, overexpression of Macf18-GFP might enhance microtubule growth rate or stabilize microtubules. This could lead to elongation of microtubule tracks and thus less pausing and higher mean speeds. In conclusion, the anterograde trafficking speed of DCVs is approximately 50% faster than retrograde transport *in vivo*.

MacF18-GFP allowed us to monitor axonal microtubule dynamics in the cerebral cortex *in vivo*. We found a unidirectional polarity of axonal movement of microtubules *in vivo* with a comet run length of around 10 μm. In comparison to developing neurons, the number of moving comets/unit of axon length is relatively small^30^. The mean speed of comet growth was 0.12 μm/s, in agreement with findings in spinal cord of anesthetized mice of 0.147 μm/s^31^ and 0.09 μm/s in dendrites of cortical neurons^24^.

The newly discovered reduction in transport speed and longer pausing times at axonal *en-passant* boutons may be a mechanism to increase the availability of DCVs at presynaptic sites^13^, consistent with the notion that these sites accumulate the protein complexes that drive exocytosis and are the main locations of DCV capture and release^26^. The mechanism that reduces DCV speed remains unclear. Several mutually non-exclusive possibilities are plausible: (1) Axonal *en-passant* boutons are small compartments packed with proteins, mitochondria and synaptic vesicles. Molecular (or organelle-) crowding could contribute to slowing of DCV transport through these structures, as indicated by diffusion models in boutons^32^. (2) DCVs are mainly transported via microtubule-mediated transport mechanisms^25^. The slowing could be caused by shorter microtubule tracks in and around boutons, causing more frequent switches between tracks, in turn leading to an effective slowing and more frequent directional changes. (3) Microtubule-mediated transport is regulated through posttranslational modifications of tubulins^33^, or the motor proteins^34,35^. Synapses may accumulate kinases and other modifying enzymes to specifically affect trafficking speed at synapses. (4) DCVs are not only transported via microtubule-dependent motors, but also via actin-dependent myosin motors^23^. The presynaptic actin-myosin network may play a role in tethering DCVs to the membrane and in DCV release^36^. The highly local actin-myosin motors may also contribute to the slowing of DCV transport at presynaptic boutons. (5) Finally, fast axonal transport of DCVs is known to be inhibited by activity in a Ca^2+^-dependent manner12. Because Ca^2+^-channels are known to accumulate in presynaptic specializations, endogenous activity in the axons will preferentially increase intracellular free Ca^2+^ in boutons and slow down DCV trafficking in this area. However, the slowing effect was also observed in anesthetized mice, presumed to have a lower neuronal activity. Taken together, a number of possible mechanisms may explain the slowing of DCVs in axonal boutons to increase the availability of DCVs at presynaptic sites.

## Additional information

### Author contributions

TK and MV designed the experiments, JK and JN carried out all experiments and analyses. All authors contributed to data interpretation. JK drafted the initial manuscript which was completed by TK and all other authors. All authors approved the final version submitted for publication and agree to be accountable for all aspects of the work in ensuring that questions related to the accuracy and integrity of any part of the work are appropriately investigated and resolved. All persons designated as authors qualify for authorship, and all those who qualify are listed.

## Acknowledgements

We thank Michaela Kaiser and Claudia Kocksch for excellent technical assistance, Ruud Toonen, Jurjen Broeke, Rhode van Westen and Claudia Persoon for discussions during the ongoing project.

## Supporting Information

### Supporting Figures

Supporting Figure 1: *In vivo* imaging of labeled DCVs

Supporting Figure 2: NPY-Venus does not show overlap with markers delineating other mobile compartments

Supporting Figure 3: Thalamaocortical axons contain DCVs independently of NPY-Cherry expression

Supporting Figure 5: DCV puncta show no changes in trafficking characteristics dependent on the duration of DCV cargo reporter expression

Supporting Figure 6: Ellipsoid axonal enlargements are presynaptic boutons

### Supporting Movies

Supporting Movie 1: DCVs trafficking through thalamocortical axons imaged in an anesthetized mouse *in vivo*. Original framerate: 1 Hz, sped up to 15 Hz

Supporting Movie 2: DCVs trafficking through thalamocortical axons imaged in an awake mouse *in vivo*. Original framerate: 1 Hz, sped up to 15 Hz

Supporting Movie 3: Microtubule +-end extension imaged in an anesthetized mouse *in vivo*. Original framerate: 1 Hz, sped up to 10 Hz

Supporting Movie 4: Slowing of DCVs at presynaptic boutons imaged in an anesthetized mouse *in vivo*. Original framerate: 1 Hz, sped up to 15 Hz

## References

1. Huang, E. J. & Reichardt, L. F. Neurotrophins: roles in neuronal development and function. Annu. Rev. Neurosci. 24, 677–736 (2001).

2. Samson, A. L. et al. Tissue-type plasminogen activator: a multifaceted modulator of neurotransmission and synaptic plasticity. Neuron 50, 673–8 (2006).

3. Cohen, S. & Greenberg, M. E. Communication between the synapse and the nucleus in neuronal development, plasticity, and disease. Annu. Rev. Cell Dev. Biol. 24, 183–209 (2008).

4. van den Pol, A. N. Neuropeptide transmission in brain circuits. Neuron 76, 98–115 (2012).

5. Meyer-Lindenberg, A., Domes, G., Kirsch, P. & Heinrichs, M. Oxytocin and vasopressin in the human brain: social neuropeptides for translational medicine. Nat. Rev. Neurosci. 12, 524–538 (2011).

6. Reichmann, F. & Holzer, P. Neuropeptide Y: A stressful review. Neuropeptides (2015). :10.1016/j.npep.2015.09.008

7. Kim, T., Gondré-Lewis, M. C., Arnaoutova, I. & Loh, Y. P. Dense-Core Secretory Granule Biogenesis. Physiology 21, (2006).

8. Zahn, T. R. et al. Dense core vesicle dynamics in Caenorhabditis elegans neurons and the role of kinesin UNC-104. Traffic 5, 544–59 (2004).

9. Kwinter, D. M., Lo, K., Mafi, P. & Silverman, M. A. Dynactin regulates bidirectional transport of dense-core vesicles in the axon and dendrites of cultured hippocampal neurons. Neuroscience 162, 1001–1010 (2009).

10. Lo, K. Y., Kuzmin, A., Unger, S. M., Petersen, J. D. & Silverman, M. A. KIF1A is the primary anterograde motor protein required for the axonal transport of dense-core vesicles in cultured hippocampal neurons. Neurosci. Lett. 491, 168–73 (2011).

11. Rose, J. B. et al. Neuropeptide Y Fragments Derived from Neprilysin Processing Are Neuroprotective in a Transgenic Model of Alzheimer’s Disease. 29, 1115–1125 (2009).

12. de Wit, J., Toonen, R. F., Verhaagen, J. & Verhage, M. Vesicular Trafficking of Semaphorin 3A is Activity-Dependent and Differs Between Axons and Dendrites. Traffic 7, 1060–1077 (2006).

13. Wong, M. Y. et al. Neuropeptide delivery to synapses by long-range vesicle circulation and sporadic capture. Cell 148, 1029–38 (2012).

14. van de Bospoort, R. et al. Munc13 controls the location and efficiency of dense-core vesicle release in neurons. J Cell Biol 199, 883–891 (2012).

15. Farina, M. et al. CAPS-1 promotes fusion competence of stationary dense-core vesicles in presynaptic terminals of mammalian neurons. Elife 4, (2015).

16. Hartmann, M., Heumann, R. & Lessmann, V. Synaptic secretion of BDNF after high-frequency stimulation of glutamatergic synapses. EMBO J. 20, 5887–97 (2001).

17. Gumy, L. F. et al. MAP2 Defines a Pre-axonal Filtering Zone to Regulate KIF1- versus KIF5-Dependent Cargo Transport in Sensory Neurons. Neuron 94, 347–362.e7 (2017).

18. Shakiryanova, D., Tully, A. & Levitan, E. S. Activity-dependent synaptic capture of transiting peptidergic vesicles. Nat. Neurosci. 9, 896–900 (2006).

19. de Wit, J., Toonen, R. F. & Verhage, M. Matrix-Dependent Local Retention of Secretory Vesicle Cargo in Cortical Neurons. J. Neurosci. 29, 23–37 (2009).

20. Berbel, P. & Innocenti, G. M. The development of the corpus callosum in cats: a light- and electron-microscopic study. J. Comp. Neurol. 276, 132–156 (1988).

21. Holtmaat, A. et al. Long-term, high-resolution imaging in the mouse neocortex through a chronic cranial window. Nat. Protoc. 4, 1128–44 (2009).

22. Taraska, J. W., Perrais, D., Ohara-Imaizumi, M., Nagamatsu, S. & Almers, W. Secretory granules are recaptured largely intact after stimulated exocytosis in cultured endocrine cells. Proc. Natl. Acad. Sci. U. S. A. 100, 2070–5 (2003).

23. Bittins, C. M., Eichler, T. W., Hammer, J. A. & Gerdes, H.-H. Dominant-negative myosin Va impairs retrograde but not anterograde axonal transport of large dense core vesicles. Cell. Mol. Neurobiol. 30, 369–79 (2010).

24. Yau, K. W. et al. Dendrites In Vitro and In Vivo Contain Microtubules of Opposite Polarity and Axon Formation Correlates with Uniform Plus-End-Out Microtubule Orientation. J. Neurosci. 36, 1071–85 (2016).

25. Barkus, R. V, Klyachko, O., Horiuchi, D., Dickson, B. J. & Saxton, W. M. Identification of an axonal kinesin-3 motor for fast anterograde vesicle transport that facilitates retrograde transport of neuropeptides. Mol. Biol. Cell 19, 274–83 (2008).

26. Torrealba, F. & Carrasco, M. A. A review on electron microscopy and neurotransmitter systems. Brain Res. Brain Res. Rev. 47, 5–17 (2004).

27. del Castillo, U., Winding, M., Lu, W. & Gelfand, V. I. Interplay between kinesin-1 and cortical dynein during axonal outgrowth and microtubule organization in Drosophila neurons. Elife 4, e10140 (2015).

28. Maday, S., Twelvetrees, A. E., Moughamian, A. J. & Holzbaur, E. L. F. Axonal Transport: Cargo-Specific Mechanisms of Motility and Regulation. Neuron (2014). 10.1016/j.neuron.2014.10.019

29. Komarova, Y. et al. EB1 and EB3 control CLIP dissociation from the ends of growing microtubules. Mol. Biol. Cell 16, 5334–45 (2005).

30. Conde, C. & Cáceres, A. Microtubule assembly, organization and dynamics in axons and dendrites. Nat. Rev. Neurosci. 10, 319–332 (2009).

31. Kleele, T. et al. An assay to image neuronal microtubule dynamics in mice. Nat. Commun. 5, 4827 (2014).

32. Wragg, R. T. et al. Synaptic activity regulates the abundance and binding of complexin. Biophys. J. 108, 1318–29 (2015).

33. Schlager, M. A. et al. Basic mechanisms for recognition and transport of synaptic cargos. Mol. Brain 2, 25 (2009).

34. Akhmanova, A. & Hammer, J. A. Linking molecular motors to membrane cargo. Curr. Opin. Cell Biol. 22, 479–487 (2010).

35. Hirokawa, N., Niwa, S. & Tanaka, Y. Molecular Motors in Neurons: Transport Mechanisms and Roles in Brain Function, Development, and Disease. Neuron 68, 610–638 (2010).

36. Papadopulos, A. et al. Activity-driven relaxation of the cortical actomyosin II network synchronizes Munc18-1-dependent neurosecretory vesicle docking. Nat. Commun. 6, 6297 (2015).

37. Schwenger, D. B. & Kuner, T. Acute genetic perturbation of exocyst function in the rat calyx of Held impedes structural maturation, but spares synaptic transmission. Eur. J. Neurosci. 32, 974–984 (2010).

38. Wimmer, V., Nevian, T. & Kuner, T. Targeted in vivo expression of proteins in the calyx of Held. Pfl�gers Arch. - Eur. J. Physiol. 449, 319–333 (2004).

39. Denk, W. et al. Anatomical and functional imaging of neurons using 2-photon laser scanning microscopy. J. Neurosci. Methods 54, 151–162 (1994).

40. Preibisch, S., Saalfeld, S., Schindelin, J. & Tomancak, P. Software for bead-based registration of selective plane illumination microscopy data. Nat. Methods 7, 418–419 (2010).

41. Tinevez, J.-Y. et al. TrackMate: An open and extensible platform for single-particle tracking. Methods (2016). doi:10.1016/j.ymeth.2016.09.016

42. Seitz, A. & Surrey, T. Processive movement of single kinesins on crowded microtubules visualized using quantum dots. EMBO J. 25, 267–77 (2006).

